# Cell Instance Segmentation Via Multi-Scale Non-Local Correlation

**DOI:** 10.1101/2023.01.24.525387

**Authors:** Bin Duan, Jianfeng Cao, Wei Wang, Dawen Cai, Yan Yan

## Abstract

For cell instance segmentation on Electron Microscopy (EM) images, state-of-the-art methods either conduct pixel-wise classification or follow a detection and segmentation manner. However, both approaches suffer from the enormous cell instances of EM images where cells are tightly close to each other and show inconsistent morphological properties and/or homogeneous appearances. This fact can easily lead to over-segmentation and under-segmentation problems for model prediction, *i.e*., falsely splitting and merging adjacent instances. In this paper, we propose a novel approach incorporating non-local correlation in the embedding space to make pixel features distinct or similar to their neighbors and thus address the over- and under-segmentation problems. We perform experiments on five different EM datasets where our proposed method yields better results than several strong baselines. More importantly, by using non-local correlation, we observe fewer false separations within one cell and fewer false fusions between cells.

## 1. INTRODUCTION

Biomedical image analysis, such as cell tracking [1], neuroanatomy reconstruction [2], depends heavily on the quality of cell instance segmentation. Instance segmentation [3, 4, 5, 6, 7] targets assembling pixels that belong to the same category and simultaneously separating individual objects from their same-category neighbors. This demand for segmentation makes it more difficult since many cells are similar and homogeneous in the biomedical images, further leading to inseparable cells for CNN models.

A well-adopted approach of instance segmentation for biomedical images is based on pixel-wise classification, such as [8, 9, 10, 11, 12, 13, 9, 14, 15, 16, 17] to outline every instance by its boundary map prediction. This approach is generally well-performed, but it can easily fail to produce a good segmentation due to a small number of misclassified pixels, such as in Figure 1. A few misclassified pixels in (c) cause an over-segmentation problem of the cell. Besides the over-segmentation problem, under-segmentation can be induced by false fused prediction of two adjacent cells, reducing the total number of instances by 1. Therefore, we demand a more continuous and compact boundary map prediction to achieve a reliable instance segmentation result.

**Fig. 1:**
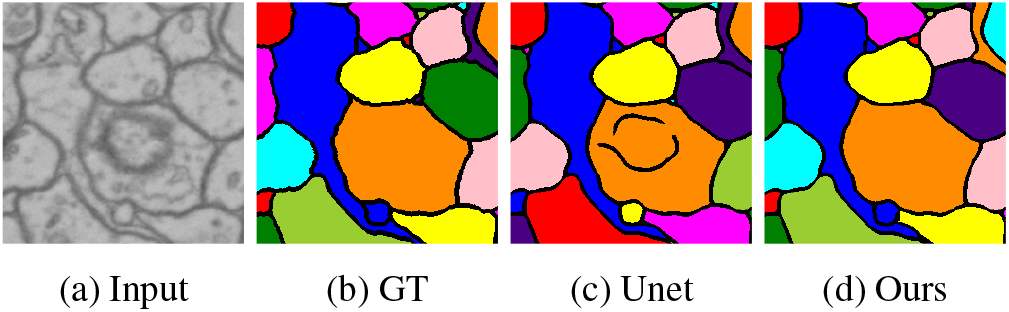
Illustration of the over-segmentation problem. GT: ground truth segmentation. The model should overlook either artifacts or nuclei within the neuron cell for better segmentation. Compared with Unet, our non-local correlation method correctly removes the artifact and produces a better result.

Another prevalent type of approach follows the detect-and-segment pipeline. The most known method, Mask-RCNN [15], first generates many region proposals and then the model segments for each proposal. Object detection requires non-maximum suppression (NMS) to remove duplicate proposals. However, for biomedical images, especially when objects have irregular shapes and inconsistent sizes, Mask-RCNN becomes problematic because there are high chances of overlapping bounding boxes of two objects, eliminating one among two authentic predictions. Hence, Mask-RCNN tends to fail to predict elongated cell structures.

In this work, we propose a novel approach utilizing the non-local correlation to learn discriminative features to obtain an accurate boundary map prediction. The reason for over-segmentation and under-segmentation is that the model falsely identifies the similarity or dissimilarity between the pixels. For example, as in Figure 1, an oracle model should recognize the artifacts by inferring from the surrounding pixels. There are studies about global [18] and local constraints [19] for assisting the model in identifying this similarity and dissimilarity. However, these methods [20, 21, 22, 23] implement sophisticated loss functions to supervise the learning of the model, which makes it hard to train. Instead of designing a new type of loss, we only use the cross-entropy loss to train a model with a fast convergence rate. Our approach enhances the ability of the model by incorporating neighboring information at the level of the feature map. Since our method operates at the feature map level, we named it *non-local correlation*. The ultimate goal of our non-local correlation is to make pixels’ features in one location either similar or unique from other locations in the embedding.

To evaluate the efficacy of our method, we compare our method with several strong baselines in five different EM datasets. Overall, our method yields comparable or better results on these datasets. By investigating the experimental results, our approach significantly reduces the extent of over-segmentation and under-segmentation problems, thus providing a higher quality of instance segmentation results. Overall, we summarize our contributions as three-fold: i) We first propose a novel non-local correlation in the EM segmentation task to reduce segmentation errors; ii) We propose a multi-scale representation to capture the correlation among different scales; iii) Extensive experimental results on five EM datasets verifies the efficacy of both non-local mechanism and multi-scale strategy.

## 2. APPROACH

Our model, shown in Fig. 2, adopts encoder-decoder architecture and outputs a boundary probability map whose value is between 0 and 1. The image encoder and correlation encoder are structurally identical but functionally different. The feature out of the correlation encoder goes through self-correlation calculation and correlation look-up and then input into the correlation bottleneck with the image feature. We then input the bottleneck output into the decoder and skip-connected the image encoder and the decoder between every feature map level except the last one to obtain the final prediction for the boundary map.

**Fig. 2:**
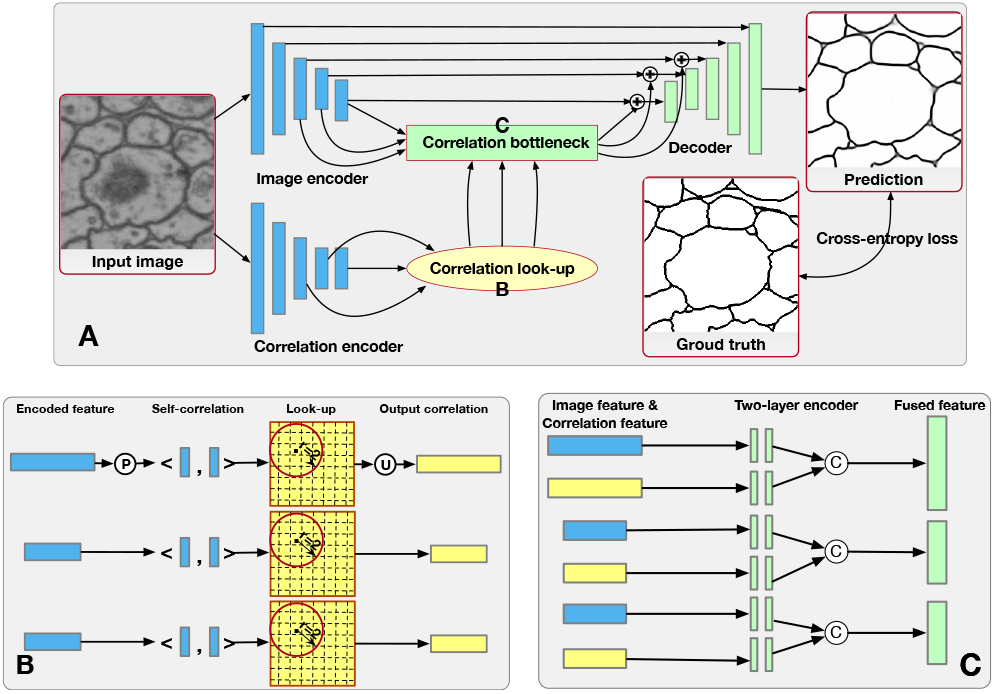
**A**. An overview of our pipeline. The input image is fed into two encoders to generate image features and correlation features. Next, we process the multi-scale correlation features with the non-local mechanism to obtain the final correlations in **B**. The correlation bottleneck **C** takes the final multi-scale correlations and the multi-scale image features and output representations as the input to the decoder to produce the boundary map prediction. 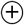, ⓒ correspondingly denote element-wise addition and concatenation. ⓟ and ⓤ denote global average pooling and upsampling, respectively.

### 2.1. Non-local correlation

Our non-local correlation operates at the feature map level, measuring the similarity between every pair of locations. Starting from an input image *x*, suppose the feature map out of the image encoder 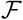 is 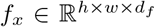, and the one output from the correlation encoder 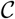 is 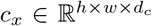, where *h, w, d* are the height, width, depths, respectively.

#### Self-correlation

Self-correlation reflects the similarity of every pair of locations within the feature map. We calculate it by the dot-product of feature *c_x_* and its transpose 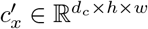, resulting in correlation representation 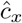

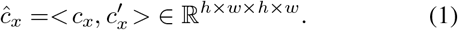

#### Correlation look-up

To index the correlation volume 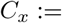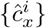, where *i* is the height of the correlation pyramid, we first initialize a grid circle with radius *r* around every location *p* for correlation feature *c_x_*, therefore the points inside the circle should be

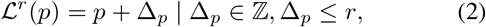

where *p* is the current location, Δ_*p*_ can be any integers between −*r* and +*r*. Next, the non-local correlation is gathered by indexing *c_x_* using 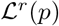, while given the mapping function *g* from coordinates to partial features of correlation representation *c_x_*, we obtain the final correlation of *c_x_* as

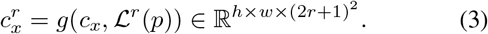

The depth of 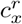 is (2*r* + 1)^2^ since the total number of points is the multiplication of the height and the width of the look-up area, which are both 2*r* plus the center itself.

#### Correlation Bottleneck output

We first embed the non-local correlation 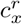 and the image feature *f*_*x*_ into the same depth, separately. Next, we concatenate the embedded feature and input it to the final bottleneck convolution to produce a representation of the same size as *f*_*x*_, used in the decoder to generate the final boundary probability map.

### 2.2. Multi-scale correlation representation

For the multi-scale representation calculation, we input multiscale image features 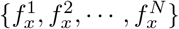 and the correlation map after correlation look-up operation into the correlation bottleneck where *N* is the maximum scale. For each pair of 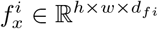 and 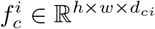, where *i* ∈ [1, *N*]. Each feature will pass a corresponding two-layer encoder and be encoded into the same dimension as the final fused feature is

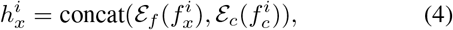

where 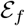 and 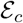 are two-layer encoders for the image feature and correlation feature, respectively, and 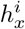 and the same size as the image feature 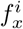.

Then, the generated multi-scale features {*h*^*i*^} | *i* ∈ [1, *N*] are added to the corresponding initial image features as the fused feature 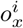

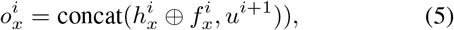

where ⊕ denotes the element-wise addition, and *U* denotes Upsampling operation. This operation first sums up correlation 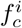 and image feature 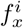 to the fused feature 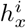. It then concatenates the fused feature *h*^*i*^ with the upsampling feature *u*^*i*+1^ of the first below layer 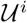 of the current level to generate the final decoder feature of current level in the decoder. During our experiments, we find the best number of the multiscale representation is three, as shown in Fig. 2.

## 3. EXPERIMENTS

### Datasets

We compare our method with other methods in five different EM datasets. **CREMI**: [24] This dataset consists of three different volumes, *i.e*., CREMI A, B, and C, which are taken from adult fruit flies, each of size 1250 × 1250 × 125 pixels, 5 × 5 × 5*μ*m^3^ of physical size. It is worth noting that these three datasets are remarkably different in appearance, especially for CREMI B and CREMI C, where cells have obscure boundaries and significant variations in size and shape. For each dataset, the first 50 slices in the *z* dimension are used in testing, and the last 75 in training. **FIB25**: [25] is of size 520 × 520 × 520, 8 × 8 × 8nm^3^ of physical size. It is challenging since it has plenty of small instances in every slice. We used the first 100 slices for testing and the rest for training the baseline model. **SNEMI3D**: [26] This image size of SNEMI3D is 1024 × 1024 × 100, 3 × 3 × 30nm^3^ of physical size. This dataset is taken from a mouse brain and comprises neuron cells that vary in shape and size. We used the first 30 slices and the last 70 to test/train the model.

### Metrics

We used two metrics for validation Arand Rand Error (ARAND) and Variation of Information (VOI). ARAND is the error version of the Adjusted Rand Index [27]. One perfect matching between prediction and ground truth has an ARAND of **0**. Conversely, ARAND of **100** indicates that nothing is matched. VOI [28, 29] measures the conditional entropy (H) between the predicted *S* and the GT segmentation *S*^⋆^, it has two components: VOI_split_ = 100 H(S^⋆^|S), VOI_merge_ = 100 × H(S|S^⋆^). VOI_split_ can be interpreted as the amount of over-segmentation, and VOI_merge_ as the amount of under-segmentation.

### Discussion of results

We show some qualitative results in Fig. 3 and quantitative results in Table 1. Overall, we observe that the competitions among different methods are tight. This relatively close performance largely owes to the strong baseline of the encoder-decoder structure. Despite the close performance, we specifically find that our non-local method improves more on VOI metrics. These performance gains certify our assumption that using the non-local mechanism can reduce false separations and fusions among neighboring cells.

**Table 1:**
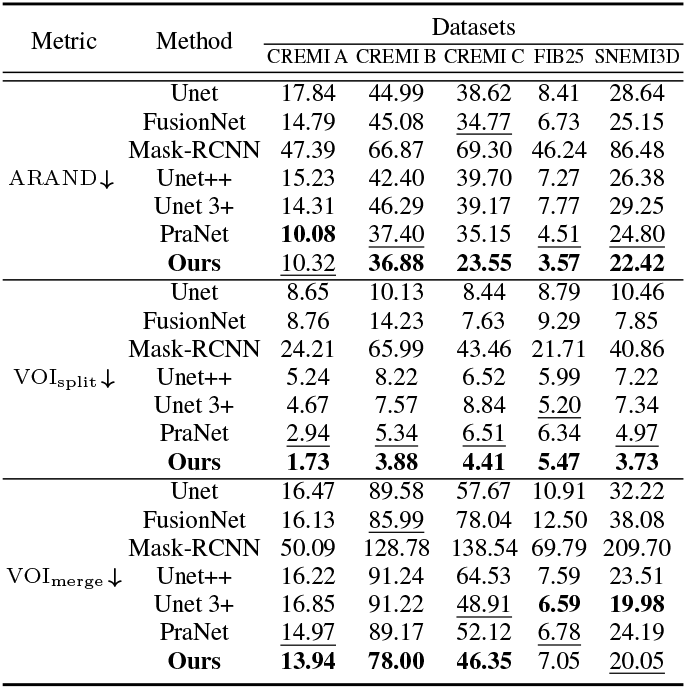
Quantitative comparisons with different methods on five datasets.

**Fig. 3:**
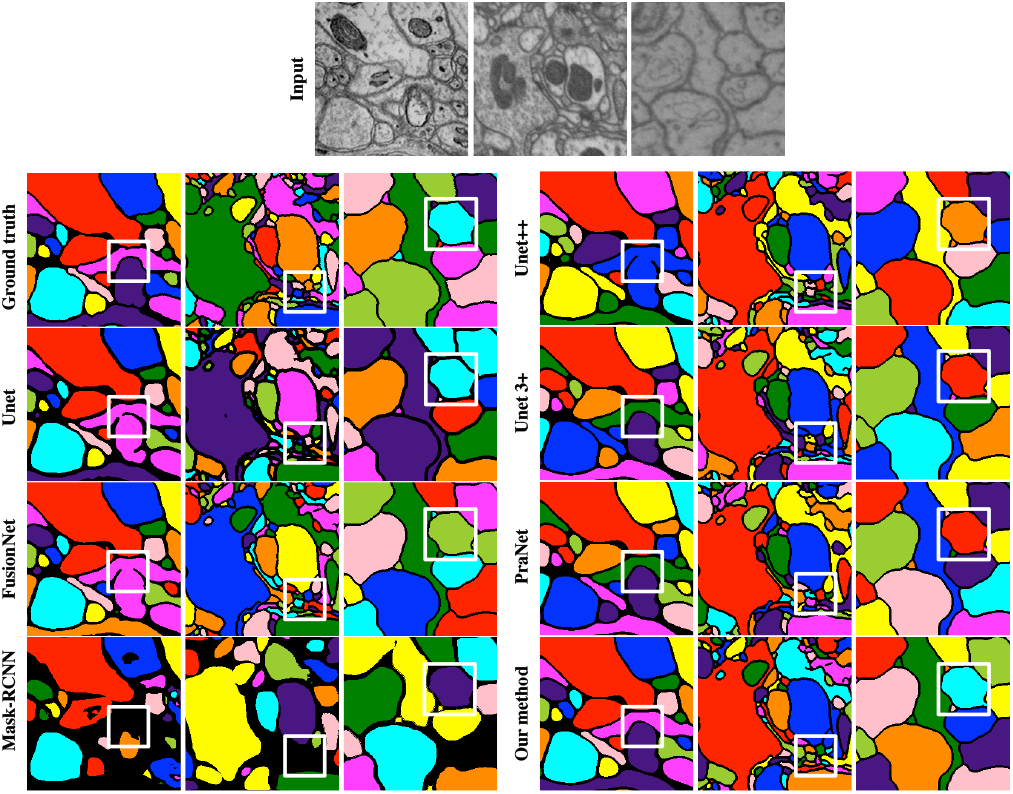
Qualitative results of different methods on different datasets: The first row are raw images: SNEMI3D (left), FIB25 (Middle), CREMI A (right). We can witness that a small boundary error can lead to over-segmentation and under-segmentation problems, as shown in the white bounding boxes. Overall, our method produces fewer false merging of adjacent cells than other methods. Mask-RCNN suffers inferior predictions due to the cells’ irregular shapes.

Specifically, the performance of all models in CREMI B and CREMI C is worse than in the other three datasets. The difficulty of the corresponding datasets induces this performance degeneration due to the varying cell size and undesired organelle boundaries. However, among all baselines, our non-local method is more resilient to cases where baseline methods fail. This robustness verifies that introducing a non-local correlation mechanism can erase some negative impact on the dataset, resulting in better segmentation. In addition, the inferior performance of Mask-RCNN can be attributed to the irregular and diverse morphology of the cells, making the region proposal less appropriate, which is a vital step in Mask-RCNN. Therefore, Mask-RCNN needs to handle flawed region proposals, which further leads to unfaithful and fragmented predictions. This tendency is apparent when long and irregular cells are segmented. Overall, our method performs better than the other baselines, leveraging the compactness of the cells with fewer over-or-under segmentations.

### Error map of segmentations

Compared with PraNet, our method achieves slightly better results in terms of boundary prediction, associated with fewer false fusions of cells (undersegmentation) and more accurate prediction within individual cells (less over-segmentation). We show the detailed error map of a problematic input image constructed between the segmentation result of PraNet and our method in Fig. 4. To sum up, our approach to learning non-local correlation significantly helps to reduce over-segmentation.

**Fig. 4:**
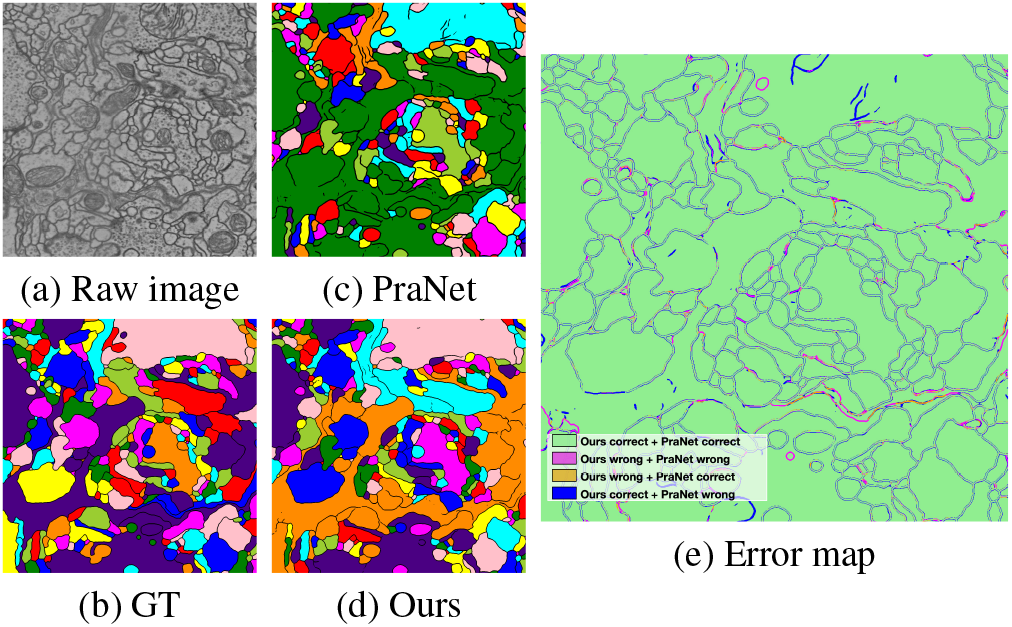
Error map between the state-of-the-art and our method (color indicated: see the color bar inside **e**). Compared to PraNet, our method is more robust in cases such as organelles with a clear boundary but should be overlooked to obtain a better segmentation. Note that the images are re-scaled for visualization purposes. Best view zoomed.

### Ablation study on numbers of multi-scale representation

We also conduct an ablation study on the number of multiscale representations for our method. The results are shown in Table 2. Due to the existence of a memory-consuming self-correlation operation, we limit our scale in {1, 2, 3}, where one means we only use the last layer of the encoder, 2 represents the last two layers of the encoder, 3 represents the last three layers of the encoder. If a full-scale correlation is urged, we would have to downsample the larger layers and upsampling back to their original size after calculating self-correlation to address the memory issue. Despite this minor limitation on the number of scales, we can observe a reasonable gain from our multi-scale correlation design.

**Table 2:** Ablation studies on different numbers of the multi-scale representation on different datasets. we fix the radius parameter at one and conduct all the experiments.

| Datasets | Setting | Metrics |  |  |
| --- | --- | --- | --- | --- |
|  |  | ARAND↓ | VOI<sub>split</sub>↓ | VOI<sub>merge</sub>↓ |
| CREMI A | Scale 1 | 12.99 | 3.00 | 14.84 |
|  | Scale 2 | 12.17 | 2.12 | 14.81 |
|  | Scale 3 | **10.32** | **1.73** | **13.94** |
| CREMI B | Scale 1 | 38.84 | 51.30 | 82.00 |
|  | Scale 2 | 37.68 | 4.95 | 81.99 |
|  | Scale 3 | **36.88** | **3.88** | **78.00** |
| CREMI C | Scale 1 | 24.51 | 5.80 | 49.63 |
|  | Scale 2 | 24.50 | 5.01 | 48.12 |
|  | Scale 3 | **23.55** | **4.41** | **46.35** |
| FIB25 | Scale 1 | 6.08 | **5.38** | 8.12 |
|  | Scale 2 | 5.10 | 6.12 | 8.49 |
|  | Scale 3 | **3.57** | 5.47 | **7.05** |
| SNEMI3D | Scale 1 | 24.01 | 5.21 | 25.32 |
|  | Scale 2 | 22.89 | 4.32 | 24.20 |
|  | Scale 3 | **22.42** | **3.73** | **20.05** |

### Ablation study on radius

To evaluate the sensitivity of the parameter radius *r*, we test our method with different radii. The result is shown in Table 3. Best performances are achieved in different radii in different datasets. Recall that our non-local correlation is learned from embedding space, which gathers information from other locations for every location to make it either heterogeneous or homogeneous. For datasets, typically CREMI A with fewer variations on the appearance of cells (shape, size, or texture), small radii are efficient enough to capture the correlation within the encoded features since larger radii may lead to faithless correlations. For datasets that are significantly dissimilar in cells, we have to choose larger radii, such as SNEMI3D. Finding an optimal radius might require some fine-tuning experiments. We fixed the radius as 1 in comparison with baseline methods.

**Table 3:** Ablation studies on different radii, where the number for multi-scale representation is 3 for these experiments.

| Metric | Setting | Datasets |  |  |  |  |
| --- | --- | --- | --- | --- | --- | --- |
|  |  | CREMI A | CREMI B | CREMI C | FIB25 | SNEMI3D |
| ARAND↓ | Radius 1 | 10.32 | 36.88 | 23.55 | 3.57 | 22.42 |
|  | Radius 2 | **9.98** | **33.04** | 29.70 | 4.27 | **20.31** |
|  | Radius 4 | 13.31 | 39.36 | 31.97 | 3.77 | 22.95 |
|  | Radius 6 | 13.97 | 41.15 | **22.37** | **3.51** | **16.98** |
|  | Radius 8 | 13.23 | 42.40 | 29.15 | **3.51** | 21.80 |
| VOI<sub>split</sub>↓ | Radius 1 | **1.73** | **3.88** | **4.41** | 5.47 | **3.73** |
|  | Radius 2 | 2.10 | 4.24 | 6.56 | **5.20** | 3.83 |
|  | Radius 4 | 2.44 | 5.82 | 6.05 | 5.59 | 5.22 |
|  | Radius 6 | 1.97 | 5.17 | 8.18 | **5.20** | 6.34 |
|  | Radius 8 | 1.94 | 4.54 | **5.51** | 5.34 | 6.47 |
| VOI<sub>merge</sub>↓ | Radius 1 | 13.94 | 78.00 | 46.35 | 7.05 | 20.05 |
|  | Radius 2 | **12.91** | **71.37** | 59.75 | 7.83 | 20.02 |
|  | Radius 4 | 17.22 | 90.13 | 64.84 | **6.50** | 23.51 |
|  | Radius 6 | 17.85 | 92.12 | **44.98** | 6.81 | **19.98** |
|  | Radius 8 | 16.97 | 99.57 | 57.72 | **6.78** | 23.49 |

## 4. CONCLUSION

Considering the nature of Electron Microscopy images containing many objects with irregular shapes and varied sizes, we propose to fully exploit the non-local correlation of the feature map in the embedding space to take the single location and multiple locations around it into account for making predictions. Our correlation mechanism enables the model to learn from neighbors around one location to distinguish or homogenize itself to its surroundings. We also find that Mask-RCNN may not be adequate to process such images with many instances that have irregular and inconsistent shapes since NMS tends to erase valid region proposals, resulting incomplete segmentation.

Our approach yields state-of-the-art results in five different EM datasets compared with other baselines. More importantly, as we discussed, we observe fewer false splittings within the same cell and fewer merges between cells in our predicted segmentation, which is crucial to separate cells.

## 5. COMPLIANCE WITH ETHICAL STANDARDS

Here, we propose a multi-scale non-local correlation mechanism on the feature map in the embedding space to boost the medical image representations and thus address the oversegmentation and under-segmentation problems. To best of our knowledge, our method may not raise any ethical issues.

## 6. ACKNOWLEDGEMENTS

This research is supported by NIH BRAIN initiative grants RF1MH123402 and RF1MH124611. This article solely reflects the opinions and conclusions of its authors and not the funding agents.

